# Active head motion reduction in Magnetic Resonance Imaging using tactile feedback

**DOI:** 10.1101/595777

**Authors:** Florian Krause, Caroline Benjamins, Judith Eck, Michael Luehrs, Rick van Hoof, Rainer Goebel

## Abstract

Head motion is a common problem in clinical as well as empirical (functional) Magnetic Resonance Imaging applications, as it can lead to severe artefacts that reduce image quality. The scanned individuals themselves, however, are often not aware of their head motion. The current study explored whether providing subjects with this information using tactile feedback would reduce their head motion and consequently improve image quality. In a single session that included six runs, 24 participants performed three different cognitive tasks: (1) passive viewing, (2) mental imagery, and (3) speeded responses. These tasks occurred in two different conditions: (a) with a strip of medical tape applied from one side of the MR head-coil, via the participant’s forehead, to the other side, and (b) without the medical tape being applied. Results revealed that application of medical tape to the forehead of subjects to provide tactile feedback significantly reduced both translational as well as rotational head motion. While this effect did not differ between the three cognitive tasks, there was a negative quadratic relationship between head motion with and without feedback. That is, the more head motion a subject produced without feedback, the stronger the motion reduction given the feedback. In conclusion, the here tested method provides a simple and cost-efficient way to reduce subjects’ head motion, and might be especially beneficial when extensive head motion is expected a priori.

## 1. Introduction

Head motion is a very common and considerable problem in clinical as well as empirical Magnetic Resonance Imaging (MRI) applications. Being one of the most frequent sources of artefacts, bulk head motion negatively affects the quality of the recorded images (for a review see Zaitsev, Maclaren & Herbst, 2015). For functional MRI (fMRI) recordings, the issue is usually addressed by retrospectively correcting the data with information from either the functional images themselves (Friston, Ashburner, Frith, Poline, Heather & Frackowiak, 1995; Friston, Williams, Howard, Frackowiak, & Turner 1996) or real-time motion tracking with a camera (Todd, Josephs, Callaghan, Lutti & Weiskopf, 2011; Stucht, Danishad, Schulze, Godenschweger, Zaitsev & Speck, 2015). However, computational algorithms for motion correction are known to leave residual motion-related artefacts in the data (Friston et al., 1996; Maclaren, Herbst, Speck & Zaitsev, 2013; Power, Mitra, Laumann, Snyder, Schlaggar & Petersen, 2014; Beall & Lowe, 2014) and can even induce false fMRI activations (Yakupov, Lei, Hoffmann & Speck, 2017). Therefore, other solutions aim to address the issue at the source and try to prevent head motion from occurring by immobilising the subject, for instance by fixating the subject’s head with a plaster cast head holder (Edward et al., 2000) or a bite bar (Bettinardi et al., 1991; Menon, Lim, Anderson, Johnson & Pfefferbaum, 1997). Unfortunately, these *passive* head motion reduction methods are cumbersome to set up and lead to significant discomfort, which is why they are not commonly used (Zaitev, Maclaren & Herbst, 2015).

The frequent occurrence of head motion even when subjects are explicitly told not to move, and the consequent need for methods to reduce it, suggest that subjects do not seem to be aware that they move their head during scanning. In fact, it has been demonstrated that providing them with this information visually, in real-time, significantly reduces head motion (Yang, Ross, Zhang, Stein & Yang, 2005; Greene et al., 2018). While this *active* head motion reduction method is very promising in general, the specific implementation does not come without costs. First, it is based on a rather complex technical setup, that includes the real-time analysis of head motion parameters, which might not be feasible to implement in some scanning facilities. Second, the information that is fed back to the participant needs to be superimposed on any visual experimental stimuli, which can potentially alter neural responses (Yang et al., 2005). Third, subjects need to learn to extract the visual feedback information from the display, which constitutes an additional task requiring additional cognitive resources (Sulzer et al., 2013; Krause et al., 2017).

Here, the potential benefits of an alternative, much simpler method to provide real-time head motion information to a subject, which does not suffer from the above-mentioned issues, are investigated: a strip of medical tape is applied from one side of the MR head-coil, via the subject’s forehead, to the other side (see Figure 1). In this setup, any head motion will produce a slight shift of the medical tape on the skin, giving immediate tactile feedback. While this method has been in active use over the last years by several researchers (including the authors, but also mentioned in Greene, Black & Schlaggar, 2016), to our knowledge no objective systematic investigation has taken place in order to verify and quantify the effects on motion reduction empirically. The current study addresses this lack of research. In a single session that included six runs, 24 participants performed three different cognitive tasks: (1) passive viewing, (2) mental imagery, and (3) speeded responses. These tasks occurred in two different conditions: (a) with a strip of medical tape applied from one side of the MR head-coil, via the participant’s forehead, to the other side, and (b) without the medical tape being applied. The tasks were chosen for being most representative of common fMRI paradigms with potentially different degrees of motion. For all three cognitive tasks, reduced motion was expected when the medical tape was applied.

**Figure 1.**
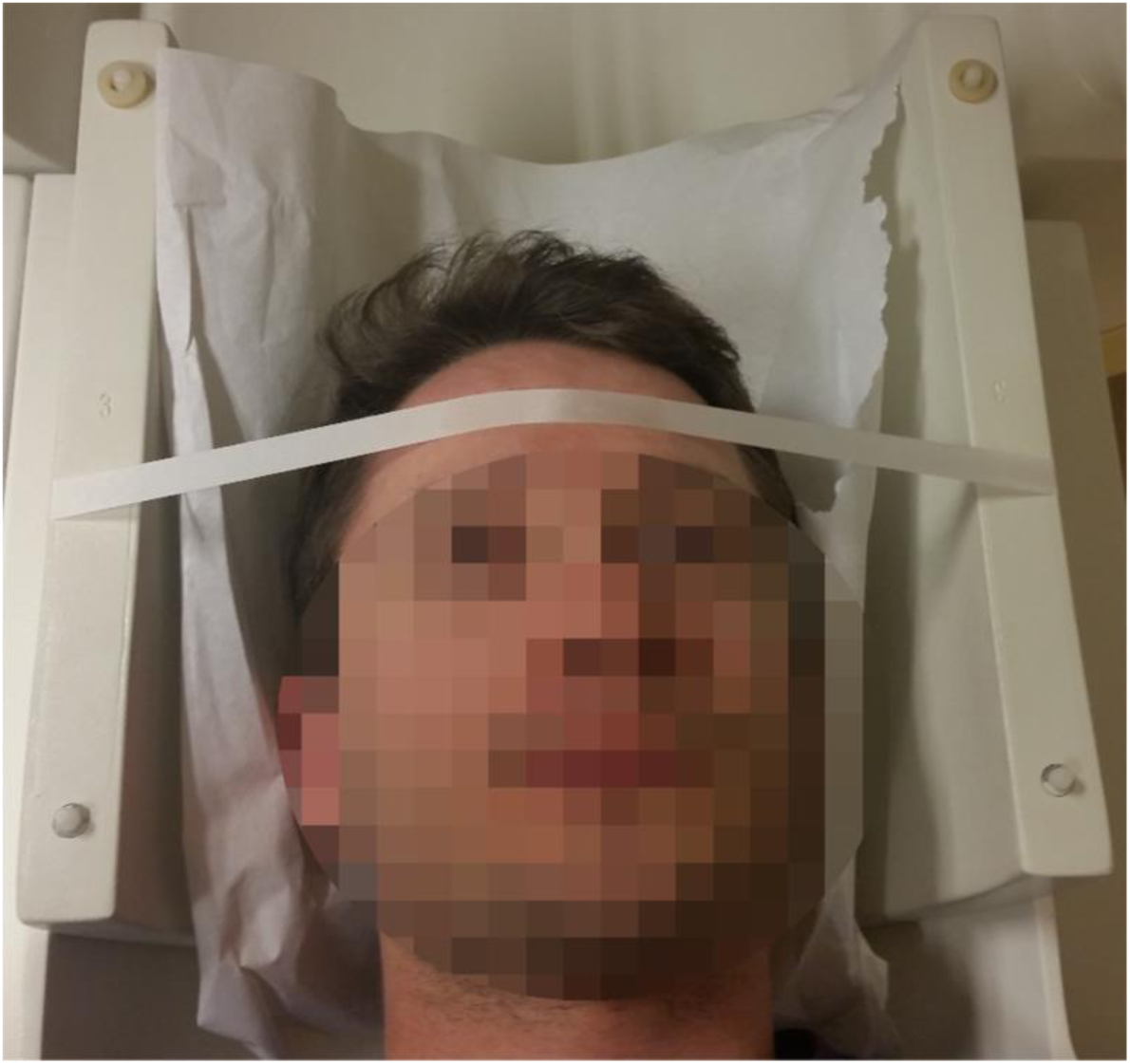
An illustration of how the medical tape was applied. For safety reasons, the picture was taken in a mock scanner and does not depict the exact same head-coil used in the current study.

## 2. Material and methods

### 2.1. Participants

Twenty-four healthy volunteers (19 females; 2 left-handed; all recruited at Maastricht University, Maastricht, the Netherlands) aged between 18 and 25 years (mean = 20.13; SD = 1.65) participated in the experiment in return for credit points. All of them had normal or corrected to normal vision and had no known neurological or psychological disorders. All volunteers had no prior fMRI experiences and have never been in an MR scanner before. The study was approved by the local ethics committee, and participants gave their written informed consent before the procedure.

### 2.2. Experimental design and procedure

Participants were engaged in a single MR session that entailed an anatomical recording followed by six functional runs. Participants were instructed to lie as still as possible throughout the entire procedure. Table 1 shows an overview of the experimental design. Each functional run consisted of 200 volumes and entailed one of three cognitive tasks: (1) passive viewing, (2) mental imagery, or (3) speeded responses. The three tasks were presented twice in two parts of the experiment, where the second part was a repetition of each task from the first part in a slightly different variant. In the passive viewing task, participants were instructed to look at a red fixation cross in the centre of the screen, while alternating blocks of 32 pictures of houses (variant 1) or objects (variant 2) and female (variant 1) or male (variant 2) faces (stimuli were identical to those described in “Photos used for FFA and LOC localization” in Kriegeskorte et al., 2003) were presented at a rate of one picture per 500 ms (leading to a block length of 16 seconds), with a rest period of 16 seconds (in which only the fixation cross was visible) in between blocks. In the mental imagery task, participants closed their eyes and were instructed to alternately rest (i.e. let their thoughts drift, but not think about anything specific) on the auditory cue “rest” and to mentally imagine to swim (variant 1) or play tennis (variant 2) on the auditory cue “swim” or “play”, respectively. Rest and imagery blocks both lasted for 16 seconds. In the speeded responses task, participants were engaged in a colour Stroop task (variant 1) or a spatial Stroop task (variant 2). In the colour Stroop task, the words “RED and “GREEN” were presented at the centre of the screen in either red or green colour, and the participants had to respond with a left button press (right index finger) if the colour of the word was green and with a right button press (right middle finger) if the colour of the word was red. In the spatial Stroop task, the words “UP” and “DOWN” were presented in either the upper or the lower half of the screen (but in the horizontal centre), and participants had to respond with a left button press (right index finger) if the word was “DOWN” and with a right button press (right middle finger) if the word was “UP”). The order and temporal spacing (between 2000 ms and 16000 ms) of trials were created randomly before the experiment and was the same for each participant.

**Table 1.** Overview of experimental design. In six runs, each participant was presented with three cognitive tasks: Passive Viewing, Mental Imagery, Speeded Responses. The three tasks were presented in two parts, with the second part being a repetition of each task from the first part in a slightly different variant: Passive 1 = Houses vs. Female Faces, Passive 2 = Objects vs. Male Faces, Imagery 1 = Mental Swimming, Imagery 2 = Mental Tennis, Response 1 = Colour Stroop, Response 2 = Spatial Stroop. In each part, medical tape was either applied (turquoise) or not (red).

|  | Part 1 |  |  | Part 2 |  |  |
| --- | --- | --- | --- | --- | --- | --- |
|  | Task 1 | Task 2 | Task 3 | Task 1 | Task 2 | Task 3 |
| Participant 01/13 | Passive 1 | Imagery 1 | Response 1 | Passive 2 | Imagery 2 | Response 2 |
| Participant 02/14 | Passive 1 | Imagery 1 | Response 1 | Passive 2 | Imagery 2 | Response 2 |
| Participant 03/15 | Passive 2 | Imagery 2 | Response 2 | Passive 1 | Imagery 1 | Response 1 |
| Participant 04/16 | Passive 2 | Imagery 2 | Response 2 | Passive 1 | Imagery 1 | Response 1 |
| Participant 05/17 | Imagery 1 | Response 1 | Passive 1 | Imagery 2 | Response 2 | Passive 2 |
| Participant 06/18 | Imagery 1 | Response 1 | Passive 1 | Imagery 2 | Response 2 | Passive 2 |
| Participant 07/19 | Imagery 2 | Response 2 | Passive 2 | Imagery 1 | Response 1 | Passive 1 |
| Participant 08/20 | Imagery 2 | Response 2 | Passive 2 | Imagery 1 | Response 1 | Passive 1 |
| Participant 09/21 | Response 1 | Passive 1 | Imagery 1 | Response 2 | Passive 2 | Imagery 2 |
| Participant 10/22 | Response 1 | Passive 1 | Imagery 1 | Response 2 | Passive 2 | Imagery 2 |
| Participant 11/23 | Response 2 | Passive 2 | Imagery 2 | Response 1 | Passive 1 | Imagery 1 |
| Participant 12/24 | Response 2 | Passive 2 | Imagery 2 | Response 1 | Passive 1 | Imagery 1 |

The rationale for having two variants of the three cognitive tasks was to make the participants believe that they were engaged in *six* different tasks, distracting them from the main manipulation in the current study: in one of the two parts a strip of medical tape (Leukopor 2.5cm; BSN medical Luxembourg Finance Holding S.à r.l., Luxembourg) was applied from one side of the MR head-coil, via the subject’s forehead, to the other side (see Figure 1). To further distract participants from the actual aim of the study, a Vitamin E supplement pill (Holland & Barrett B.V., The Netherlands) was attached to the tape and participants were told that the reason for this procedure was to better locate their head position in the scanner. Since the actual interest of the current study was to investigate the effects of the medical tape as an *active* head motion reduction method, participants needed to know that any head motion would produce a slight shift of the medical tape on their skin, giving immediate tactile feedback, and that they could use this information to fulfil the requirement of lying as still as possible. Participants were given this information in a mere side remark while applying the medical tape, in order to prevent them from realising the main aim of the study. Whether the medical tape was applied during the first or second part of the experiment was alternated from participant to participant. When the medical tape was applied in the second part, it was applied after three tasks and participants were told that this was needed for the subsequent scans. When the medical tape was applied in the first part, it was removed after three tasks and participants were told that it is no longer needed for the subsequent scans. In either case, the need to lie as still as possible was reiterated by the researcher at the beginning of each part. Being MR novices, the participants did not question either change in the setup after three tasks, and none of the participants realised the main aim and manipulation of the study (according to verbal reports in the debriefing after the experiment).

After the experiment, participants were asked to fill in a short questionnaire in which they rated the difficulty of each cognitive task and how much they thought they had moved during that task on a scale from 0 to 10.

All experimental paradigms were presented using Expyriment (Krause & Lindemann, 2014). Visual stimuli were projected onto a screen at the end of the scanner bore. Auditory cues were played back via MR-compatible in-ear headphones. Speeded manual responses were recorded using an MR-compatible response box. The order of cognitive tasks, task variants and application of medical tape were counter-balanced across participants.

### 2.3. Data acquisition

MR images were recorded on a 3-T Siemens Magnetom Prisma MR system (Siemens, Erlangen, Germany) with a 64-channel receiver head coil. High-resolution sagittal anatomical images were acquired using a T1-weighted MP-RAGE sequence with a GRAPPA acceleration factor of 2 (repetition time/echo time = 2250/2.21 ms; flip angle = 9°; field of view = 256 × 256 mm; number of slices = 192; slice thickness = 1.0 mm; in-plane resolution = 1.0 × 1.0 mm). Functional images were acquired using an echo planar T2*-weighted sequence sensitive to BOLD contrast with a GRAPPA acceleration factor of 2 (repetition time/echo time = 2000/30 ms; flip angle = 77°; field of view = 216 × 216 mm; number of slices = 35; slice thickness = 3.0 mm; in-plane resolution = 3.0 × 3.0 mm). In an attempt to match brain coverage across participants, Siemens Auto-Alignment Scout (AAScout) was applied, but the exact orientation often had to be re-adjusted manually.

### 2.4. Data analysis

#### 2.4.1 Head motion parameters

All MR images were preprocessed using *BrainVoyager* (version 20.2; Goebel, 2012). Slice timing corrected functional images were realigned to the first image of each run using trilinear detection and sinc interpolation (100 iterations), resulting in the calculation of six motion parameters (three translational, three rotational). Based on these parameters, two indices were calculated each for rotational and translational motion parameters individually. The *head displacement index* captures the absolute displacement of each volume from the initial position at the beginning of the run and was calculated as

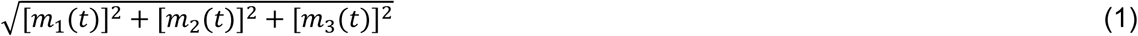

while the *head motion index* captures the relative motion from volume to volume and was calculated as

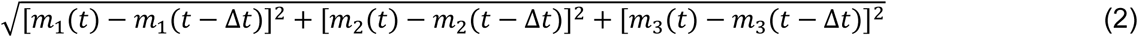

where *m*_*1*_, *m*_*2*_ and *m*_*3*_ are the three motion parameters (cf. Yang et al., 2005). In addition, framewise displacement (FD) was calculated as

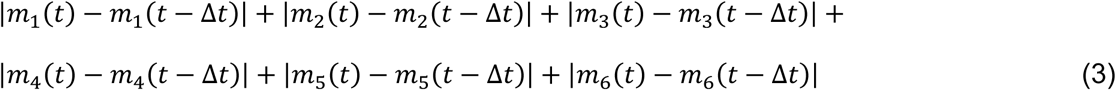

combining all six translational and rotational parameters (*m*_*1*_ – *m*_*6*_) into a single measure. Rotational displacements were converted from degrees to millimeters by calculating displacement on the surface of a sphere of radius 50 mm (c.f. Power, Barnes, Snyder, Schlaggar & Petersen., 2012).

To investigate the effect of the application of the medical tape on short term head motion (i.e. volume-to-volume motion), mean FD, as well as mean translational and rotational head motion indices per run were each entered into a separate 2 × 3 repeated measures Analysis of Variance (ANOVA) with the factors *Condition* (Tape, NoTape) and *Task* (Passive, Imagery, Responses). To investigate the effect of the application of the medical tape on long term head motion (i.e. drift), the regression coefficients of the linear regressions on translational and rotational head displacement indices per run were each entered into a separate 2 × 3 repeated measures ANOVA with the factors *Condition* (Tape, NoTape) and *Task* (Passive, Imagery, Responses). To understand the effect of the medical tape in more detail, average motion in the Tape condition (mean of all tasks per participant) was regressed on average motion in the NoTape condition with a linear and an orthogonalized quadratic term.

To investigate the effect of the application of the medical tape on between-run motion, head displacement indices were calculated in the same way as described above, but on motion parameters that resulted from an image realignment to the first image of the first run of each part (i.e. run 1, 2 and 3 were realigned to run 1 and run 4, 5 and 6 were realigned to run 4), in order to preserve information about motion between runs. For runs 2, 3, 5 and 6, the absolute difference between the first head displacement index value of the current run and the last head displacement index value of the previous run was then extracted for translation and rotation individually, and averages for both conditions (Tape, NoTape) for each participant entered a paired t-test.

All analyses based on the motion parameters were performed in Python (version 3.7.0; Python Software Foundation, 2018) and R (version 3.5.1; R Core Team, 2018) using the package ‘afex’ (version 0.22; Singmann, Bolker, Westfall & Aust, 2018).

#### 2.4.2 fMRI data

All MR images were preprocessed using BrainVoyager (version 20.2; Goebel, 2012). Anatomical images were corrected for inhomogeneities and normalised to the Montreal Neurological Institute (MNI) standard space. Slice timing corrected functional images of each run were realigned to the first image of the first run (run 1, 2 and 3) or fourth run (run 4, 5 and 6) using trilinear detection and sinc interpolation (100 iterations), resulting in the calculation of six motion parameters (three translational, three rotational). Realigned images were subsequently high-pass-filtered (linear trend removal and 2 cycles per run), co-registered to the corresponding anatomical image, and spatially smoothed with a Gaussian kernel of 4 mm FWHM.

To explore general effects of the application of the medical tape on task-independent fMRI data quality, timecourses of each run were extracted from a large number of regions throughout the brain, based on the parcellation by Gordon and colleagues (2016; 333 regions). To guarantee functional coverage in all runs of the scanned cohort, thirty regions had to be reduced in size and another thirty had to be removed from further analysis. For each of the remaining 303 regions, effects of the task and head motion were regressed out of the timecourse of each run by regressing 7 predictors (1 task, 6 motion) onto the data, resulting in a cleaned residual timecourse. For passive viewing, the task was modelled as the duration of the stimulus presentation blocks, for mental imagery, the task was modelled as the duration of the imagery blocks and for speeded responses, the task was modelled as the time between stimulus presentation and response). Head motion was modelled with the six motion parameters from image realignment. The cleaned timecourse data formed the basis for two measures: (1) the average (across all runs of all subjects) difference in temporal signal-to-noise ratio (tSNR; Welvaert & Rosseel, 2013) between conditions (Tape, NoTape) was calculated for each region, and (2) the connection strengths (Pearson correlation of cleaned timecourses) between all 303 regions were obtained, and for each condition (Tape, NoTape), the correlation between average (over all runs from all subjects) connection strength (Fisher Z-transformed Pearson correlation coefficient) and head motion (FD), was calculated (c.f. Patriot et al., 2016). All analyses on timecourse data were performed in MATLAB 2015b, The MathWorks, Inc., Natick, Massachusetts, United States, and Python (version 3.7.0; Python Software Foundation, 2018) using the package ‘SciPy’ (version 1.1.0; Jones, Oliphant, Peterson et al., 2001).

To further examine whether the head motion reduction induced by the medical tape also specifically affected task-related fMRI activations in the three cognitive tasks in the current study, a fixed-effects generalised linear model (GLM) was created for each cognitive task. Each GLM included two regressors of interest, modelling the effect of the task with and without tape, respectively (tasks were modelled as described above). In all GLMs, regressors of interest were convolved with the haemodynamic response function and the six motion parameters from image realignment were included as covariates. After accounting for serial correlations with an autoregressive AR(2) model, for each task, the contrast *Tape > NoTape* was tested at a voxel threshold of *p* < .05, corrected for multiple comparisons by means of the false discovery rate (FDR). All fMRI activation analyses were performed in BrainVoyager (version 21.2).

#### 2.4.3 Behavioural performance

To inspect whether the application of the medical tape affected behavioural performance in the speeded response task, response time data from the two variants (colour Stroop task and spatial Stroop task) was aggregated and entered into a repeated measures ANOVA with the factors *Congruency* (congruent, incongruent) and *Condition* (Tape, NoTape). All analyses based on response data were performed in R (version 3.5.1; R Core Team, 2018) using the package ‘afex’ (version 0.22; Singmann et al., 2018).

#### 2.4.4 Participant ratings

To assess participants’ subjective ratings of task difficulty and their head motion, questionnaire data were analysed. For both ratings, a separate repeated measures ANOVA with the factors *Condition* (Tape, NoTape) and *Task* (Passive, Imagery, Response) was conducted. Greenhouse-Geisser correction was applied when necessary. All analyses based on questionnaire data were performed in R (version 3.5.1; R Core Team, 2018) using the package ‘afex’ (version 0.22; Singmann et al., 2018).

#### 2.4.5 Instructed head motion

To demonstrate that the medical tape is not simply restricting motion physically, the range of possible head motion when explicitly instructed to move was explored. An additional independent participant (female, 49 years old) was asked to actively move her head as if looking in one of four directions. In two short runs (each 71 volumes) with four blocked conditions, the participant was instructed to move from a central head position to an up, down, left or right tilted head position and back (five times each) with a frequency of 0.5 Hz (paced auditorily). In one of the runs, the medical tape was applied, while in the other run it was not. For each of the two runs FD (see above) was calculated and compared with the run with the maximum head motion observed in each of the two conditions in the current study.

### 2.5. Data Availability Statement

Anonymized data and scripts for reproducing the reported analyses for head motion parameters, tSNR, functional connectivity, behavioural performance as well as participant ratings are openly available in the Open Science Framework at https://osf.io/hrnfw/.

MR images are and the related fMRI activation analyses are available on request from the corresponding author. MR images are not publicly available due to privacy or ethical restrictions.

## 3. Results

### 3.1. Head motion parameters

In line with our hypothesis, the ANOVA on mean translational head motion indices revealed a significant main effect of the factor Condition, *F*(1, 23) = 7.61, *p* < .05, *η*_*p*_^*2*^ = .25, with less translational volume-to-volume motion when the medical tape was applied (0.0171 mm), compared to when it was not (0.0241 mm). No main effect of the factor Task and no interaction between the factors Condition and Task could be observed (both *F* < 1; Figure 2 (A)). Likewise, the ANOVA on mean rotational head motion indices showed a significant main effect of the factor Condition, *F*(1, 23) = 7.35, *p* < .05, *η*_*p*_^*2*^ = .24, with less rotational volume-to-volume motion when the medical tape was applied (0.0146 deg), compared to when it was not (0.0215 deg). Again, no main effect of the factor Task (*F* = 1.37) and no interaction between the factors Condition and Task (*F* = 1.12) could be observed (Figure 2 (B)). The ANOVA on FD also revealed a significant main effect of the factor Condition, *F*(1, 23) = 8.13, *p* < .01, *η*_*p*_^*2*^ = .26, with less overall volume-to-volume motion when the medical tape was applied (0.0436 mm), compared to when it was not (0.0627 mm). No main effect of the factor Task and no interaction between the factors Condition and Task could be observed (both *F* < 1).

**Figure 2.**
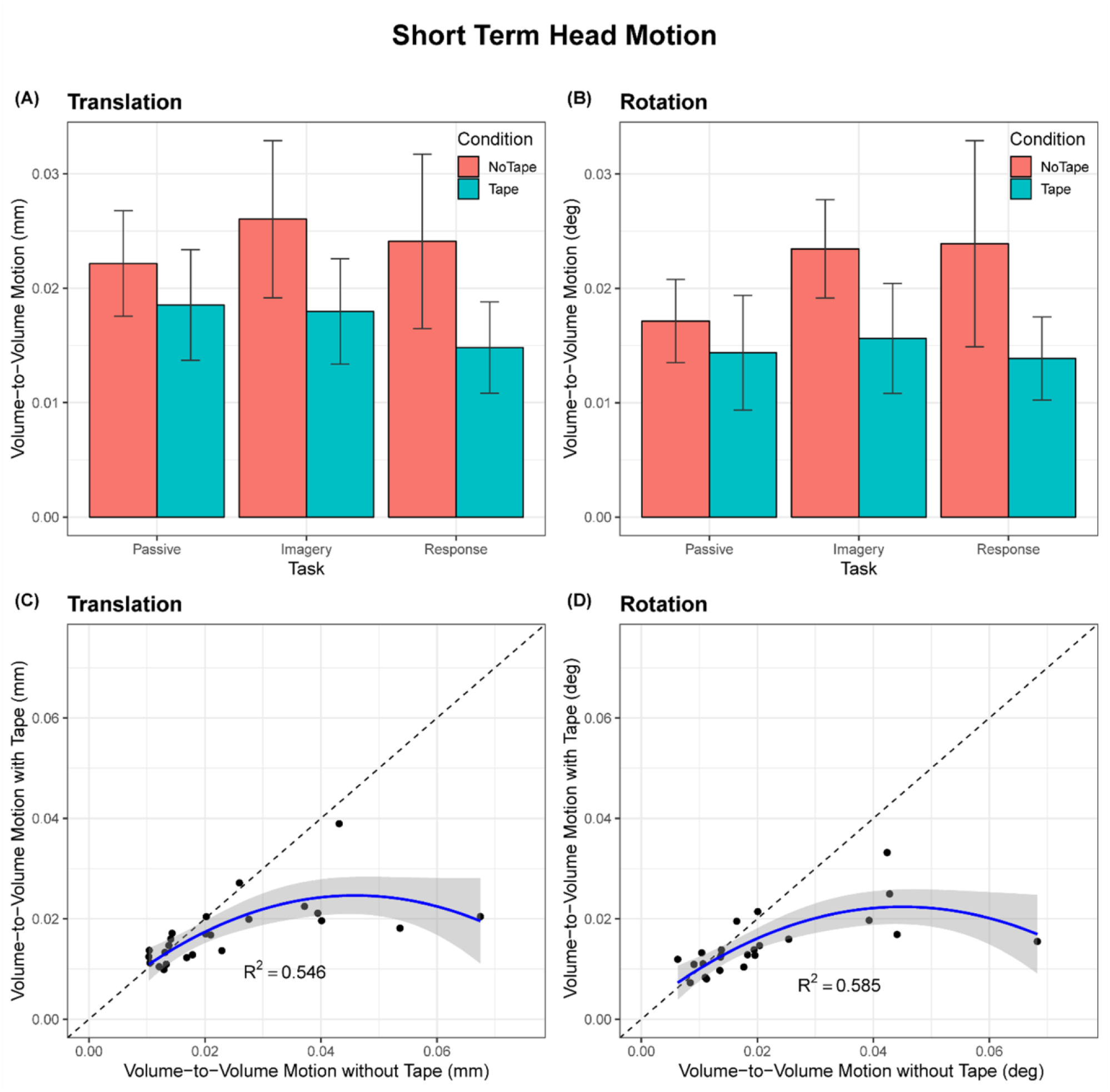
Results of analyses on short term head motion. (A) Mean translational volume-to-volume motion as a function of Task and Condition, showing a main effect of Condition. Error bars represent 95% confidence intervals for within-subject designs (Morey, 2008). (B) Mean rotational volume-to-volume motion as a function of Task and Condition, showing a main effect of condition. Error bars represent 95% confidence intervals for within-subject designs (Morey, 2008). (C) Non-linear relation between translational motion in the NoTape condition Tape condition. For short term head motion, the more motion there is without the medical tape, the stronger the advantageous effect of the medical tape. (D) Non-linear relation between rotational motion in the NoTape condition and Tape condition. For short term head motion, the more motion there is without the medical tape, the stronger the effect of the medical tape.

Furthermore, the ANOVA on the regression coefficients of the linear regressions on translational head displacement indices revealed a significant main effect of Condition, *F*(1, 23) = 5.45, *p* < .05, *η*_*p*_^*2*^ = .19, with less translational drift when the medical tape was applied (0.0023 mm/volume), compared to when it was not (0.0033 mm/volume). No main effect of the factor Task (*F* < 1) and no interaction between the factors Task and Condition (*F* = 1.08; Figure 3 (A)) were observed. Similarly, the ANOVA on the regression coefficients of the linear regressions on rotational head displacement indices showed a significant main effect of Condition, *F*(1,23) = 8.02, *p* < .01, *η*_*p*_^*2*^ = .26, with less rotational drift when the medical tape was applied (0.0021 deg/volume), compared to when it was not (0.0033 deg/volume). As in the former analyses, no main effect of the factor Task (*F* = 1.26) and no interaction between the factors Task and Condition (*F* < 1) were observed (Figure 3 (B)).

**Figure 3.**
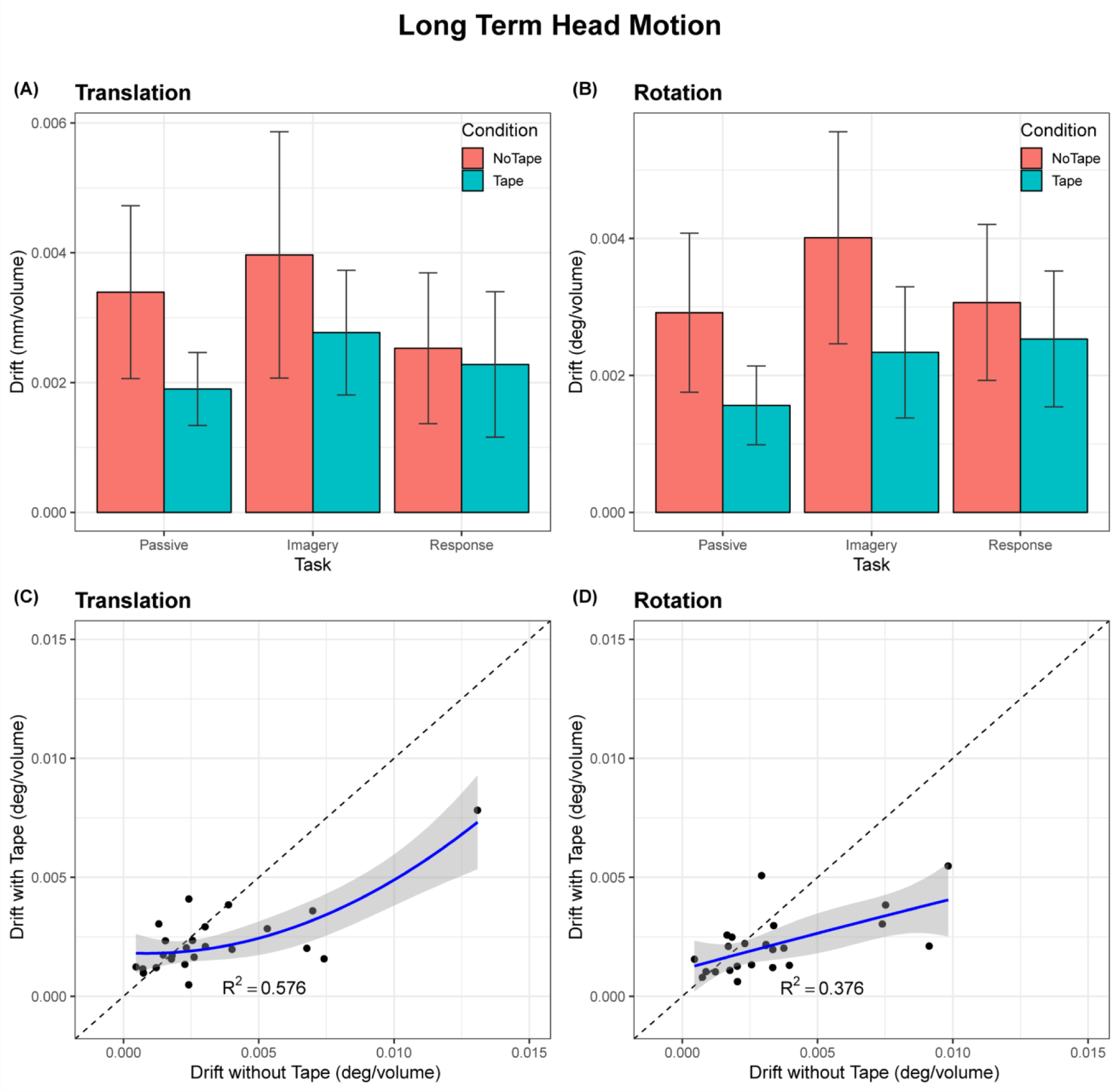
Results of analyses on long term head motion. (A) Mean translational volume-to-volume motion as a function of Task and Condition, showing a main effect of Condition. Error bars represent 95% confidence intervals for within-subject designs (Morey, 2008). (B) Mean rotational volume-to-volume motion as a function of Task and Condition, showing a main effect of Condition. Error bars represent 95% confidence intervals for within-subject designs (Morey, 2008). (C) Relation between translational motion in the NoTape condition and the Tape condition. For long term head motion, the effect of the medical tape does not scale with the amount of motion there is without the medical tape. (D) Relation between rotational motion in the NoTape condition and Tape condition. For long term head motion, the effect of the medical tape does not scale with the amount of motion there is without the medical tape.

The regressions of average motion in the NoTape condition on the average motion in the Tape condition resulted in overall significant model fits for translational volume-to-volume motion, *R*^*2*^ = .546, *p* < .001, rotational volume-to-volume motion, *R*^*2*^ = .585, *p* < .0001, FD, *R*^*2*^ = .596, *p* < .0001, translational drift, *R*^*2*^ = .576, *p* < .001, and rotational drift, *R*^*2*^ = .376, *p* < .01. Notably, the addition of the quadratic term significantly improved the model for translational volume-to-volume motion, *F*(1,21) = 8.49, *p* < .01, rotational volume-to-volume motion, *F*(1,21) = 10.83, *p* < .01, and FD, *F*(1,21) = 11.18, *p* < .01. The resulting negative non-linear relationship between the two conditions indicates that the more motion is present in a classical situation without medical tape applied, the more the medical tape helps proportionally to reduce this motion (see also Figure 2 (C—D)). The addition of the quadratic term did not improve the model for translational drift, *F*(1,21) = 2.94, *p* = 0.137, and rotational drift, *F*(1,21) = 0.05, *p* = 0.834, indicating the lack of a non-linear relationship (see also Figure 3 (C—D)).

The paired t-tests on between-run motion indicated significantly less translational head motion when the medical tape was applied (0.28 mm), compared to when it was not applied (0.68 mm), *t*(23) = 2.569, *p* < .05, *d* = 0.52, as well as significantly less rotational head motion when the medical tape was applied (0.22 deg), compared to when it was not applied (0.58 deg), *t*(23) = 2.155, *p* < .05, *d* = 0.43.

### 3.2. fMRI data

The application of the medical tape had an overall positive effect on fMRI data quality. Within the 303 sampled regions, an average increase in tSNR of 8.08 was observed (*t(302)* = 17.99, *p* < .0001). Figure 4 shows the spatial distribution of the changes in tSNR throughout the brain.

**Figure 4.**
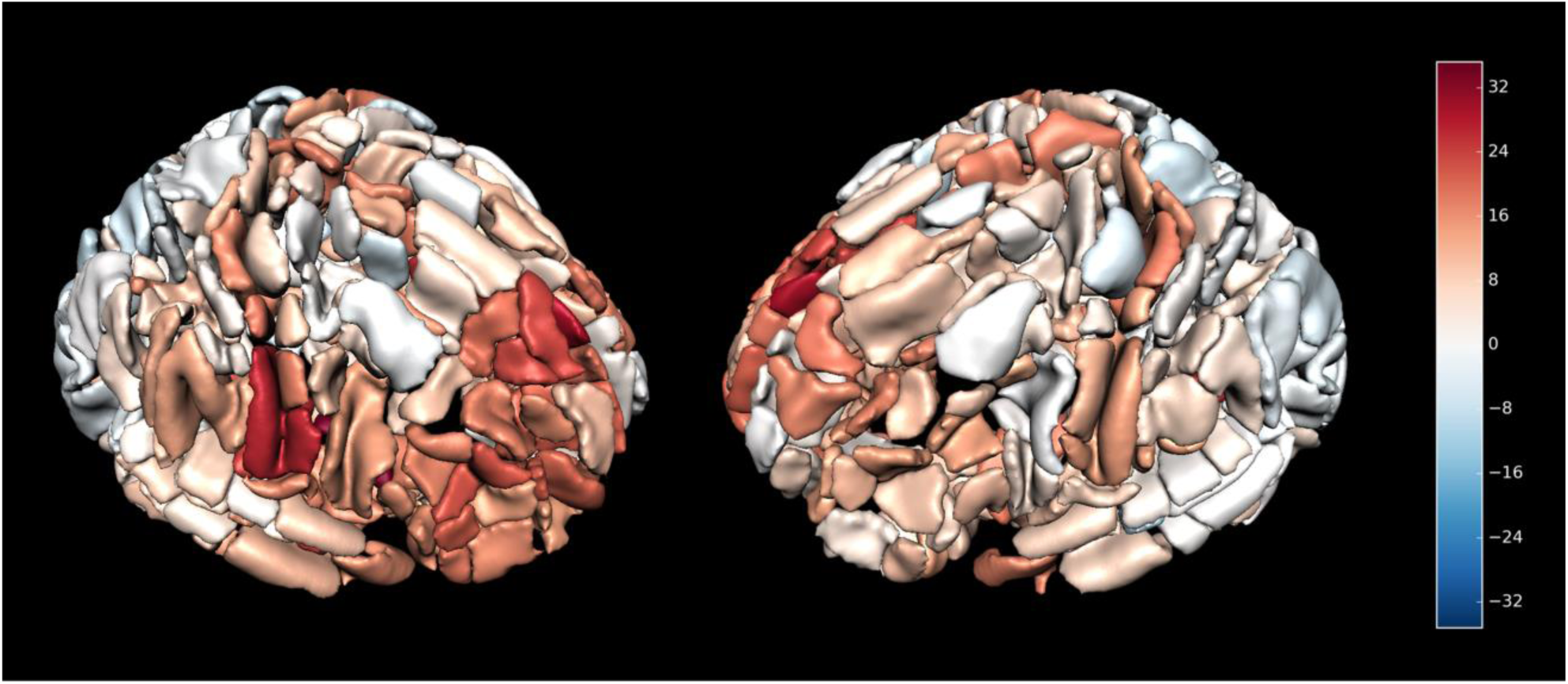
Spatial distribution of the changes in tSNR when the medical tape was applied. Overall, an increase in tSNR was observed.

Furthermore, a reduced amount of negative as well as positive correlations between head motion and connection strength was observed when the medical tape was applied, compared to when it was not applied. The strength of correlation reduction was slightly, but significantly, positively correlated with the cortical distance of the connection (*r* = 0.18, *p* < .0001). That is, the further away two regions, the more the medical tape helped to reduce the correlation between the connectivity of these regions and motion. Figure 5 shows the correlation matrices and histograms for both conditions.

**Figure 5.**
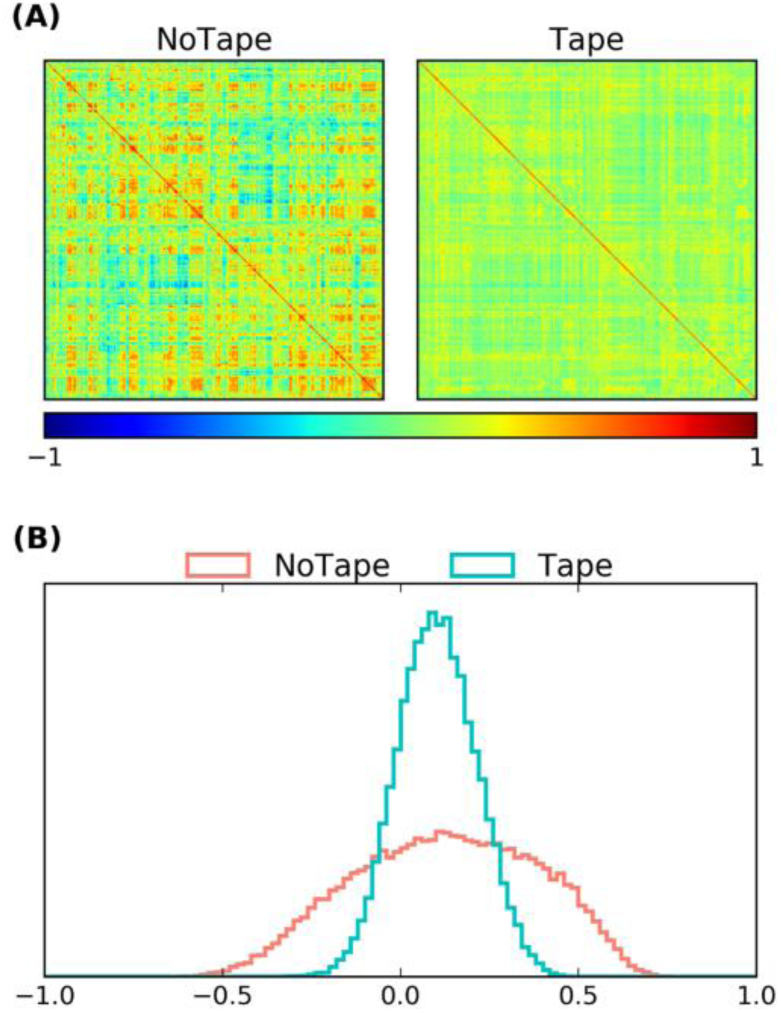
Influence of head motion on functional connectivity (A) Matrices with correlations (across all runs form all subjects) between average head motion and connection strength for 303 regions for both conditions. (B) Histograms of the correlations for both conditions. Application of the medical tape led to fewer negative and positive correlations.

Eventually, the medical tape also affected task-related fMRI activations in the tested group of participants. Figure 6 shows significant activations and deactivations of the effect of the medical tape in each of the three cognitive tasks. The application of the medical tape led to changes in bilateral early visual cortex in the passive viewing task, bilateral visual and parietal cortices in the mental imagery task, and left motor primary cortex in the speeded responses task.

**Figure 6.**
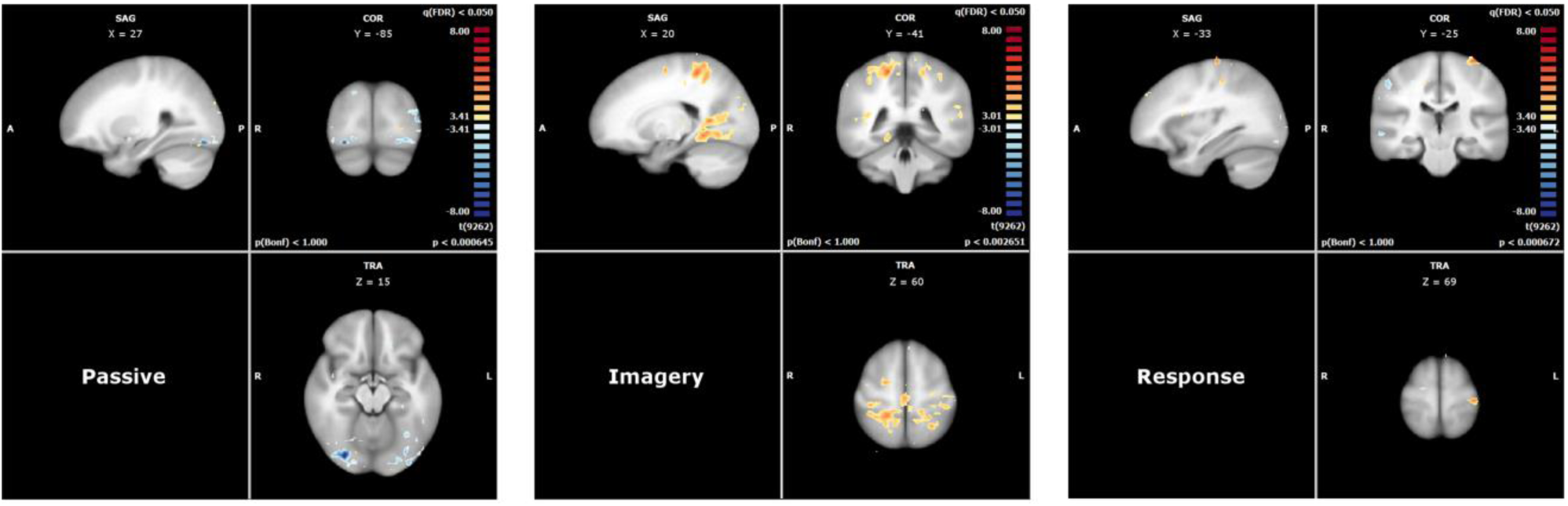
Changes in task-based fMRI activation induced by the application of the medical tape for all three cognitive tasks: bilateral early visual cortex in the passive viewing task, bilateral visual and parietal cortices in the mental imagery task, and left motor primary cortex in the speeded responses task.

### 3.3. Behavioural performance

The ANOVA on response times of the speeded response task revealed a significant main effect of Congruency, *F*(1, 23) = 70.53, *p* < .001, *η* _*p*_^*2*^ = .75, with faster responses for congruent trials (557 ms) compared to incongruent trials (609 ms), indicating a Stroop effect. No significant effects were observed for the factor Condition (*F* = 1.19) and the interaction between the factors Congruency and Condition (*F* < 1).

### 3.4. Participant ratings

The ANOVA on the participants’ ratings of perceived task difficulty in each run revealed a significant main effect of the factor Task, *F*(1.86, 42.69) = 26.34, *p* < .001, *η*_*p*_^*2*^ = .53, with a lower rating in the passive task (2.00) compared to both the imagery task (4.18; *t*(47) = 7.567, *p* < .001) and the response task (4.23; *t*(47) = 7.368, *p* < .001). No significant effects were observed for the factor Condition (*F*(1, 23) = 3.25, *p* = .08) and the interaction between the factors Condition and Task (*F* < 1). The ANOVA on the participants’ ratings of perceived head motion in each run also revealed a significant main effect of the factor Task, *F*(1.94, 44.65) = 11.7, *p* < .001, *η*_*p*_^*2*^ = .34, with a lower rating in the passive task (1.96) compared to the response task (2.46; *t*(47) = 7.920, *p* < .001) and lower rating in the response task compared to the imagery task (3.22; *t*(47) = 3.745, *p* < .001). In line with the actual head motion data, there was a significant effect of the factor Condition (*F*(1, 23) = 8.61, *p* < .01, *η*_*p*_^*2*^ = .27), with less perceived head motion when the medical tape was applied (2.35) compared to when it was not applied (2.74). No interaction between the factors Condition and Task was observed (*F* < 1).

### 3.5. Instructed head motion

Having explicitly instructed an independent additional participant to move her head, average FD was 2.48 mm during the run in which the medical tape was applied and 3.25 mm in the run in which it was not applied (Figure 7(A)). In contrast, the maximum observed average run FD in the current study was 0.27 mm without the medical tape applied (run 2 of participant 19, speeded responses) and 0.19 mm with the medical tape applied (run 4 of participant 3, passive viewing; Figure 7(B)). Those two runs also contained the maximum observed peak FD (i.e. the difference in head position between two consecutive single volumes within a run) in the current study of 12.62 mm without the medical tape applied and 5.81 mm with medical tape applied (see also Figure 7(B)). In comparison, the maximum peak FD in the additional participant explicitly instructed to move head was 14.57 mm without the tape applied and 12.57 mm with the tape (see also Figure 7(A)).

**Figure 7.**
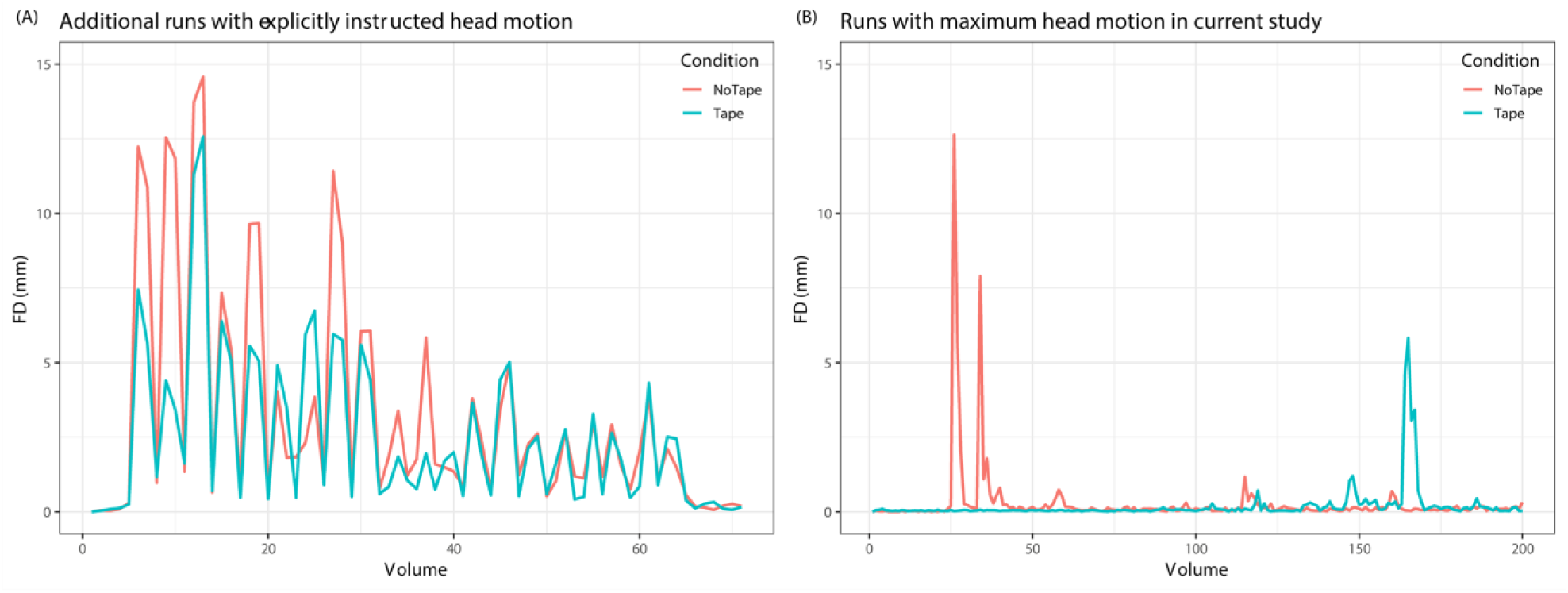
Comparison of possible head motion (when explicitly instructed) with head motion observed in the current study. (A) Runs of an independent additional participant who was explicitly instructed to move her head, once with the medical tape applied and once without. (B) Runs with maximum head motion in the current study per condition (with medical tape, without medical tape applied). The effect of the medical tape on head motion was not simply due to physical restriction.

## 4. Discussion

The current study is a first investigation of the efficacy of a simple tactile-feedback-based method for the voluntary reduction of subject head motion during MR recordings that has been employed by several researchers over the last years: a strip of medical tape that is applied from one side of the MR head-coil, via the subject’s forehead, to the other side. Our results indicate that this method significantly reduced short-term motion (i.e. volume-to-volume) as well as long-term motion (i.e. drift) in both the translational and rotational domain, making this simple tactile-feedback-based active motion reduction method a viable alternative (or addition) to other motion reduction methods.

Interestingly, the observed head motion reduction was not dependent on the particular task being performed in the scanner (passive viewing, motor imagery, speeded responses), suggesting that it can be beneficial in a large variety of MR applications, ranging from rather passive structural or resting-state recordings to more active functional experimental paradigms. Short term head motion reduction did, however, scale with the amount of head motion. That is, the more head motion an individual produced without medical tape applied, the more beneficial the application of medical tape became. This suggests that the here tested method would be especially beneficial for subject populations in which more head motion can be expected a priori (e.g. children; Greene et al., 2016; Greene et al., 2018). This becomes particularly important, when such populations are to be compared to a control population, since strong between-group differences in head motion can introduce spurious results in MRI data (Green et al., 2016). In this context, it is worth noting that even though the effect of the tape in the current study was of substantial size (*η*_*p*_^*2*^ between .18 and .25), head motion in the tested group of participants was remarkably low already without the medical tape applied. One potential explanation for this might lie in the fact that the study design incorporated multiple short functional runs (of about 6 minutes each). The frequent occurrences of restful breaks (i.e. no scanner noise, no task demands) resulting from this particular setting might have led to less discomfort- and exhaustion-related head motion than a more common long functional run of up to an hour, in which the effects of the medical tape on head motion can be expected to be even stronger.

The explicit choice of using multiple short functional runs in the current study did, however, allow for analysing head motion between subsequent experimental runs, which the medical tape also significantly reduced. This makes the application of the medical tape especially useful for real-time fMRI applications which target specific pre-defined brain regions and rely on accurate positioning information during a session with multiple measurements, such as brain-computer interfaces and neurofeedback (Weiskopf et al., 2004; Weiskopf et al., 2003). Since real-time fMRI analyses also cannot incorporate some of the more advanced and computationally demanding data-driven retrospective motion correction methods often used for offline data (e.g. Independent Component Analysis; Pruim et al., 2015), they also strongly benefit from the within-run motion reduction the medical tape provides.

The negative effects of head motion on (f)MRI data are generally so well acknowledged, with many different proposed methods that attempt to correct for it in retrospect (see Zaitsev, Maclaren & Herbst, 2015 for a review), that it should make the benefit of the here presented significant prospective motion reduction rather obvious: it is always preferable to prevent head motion from happening, rather than trying to correct for it afterwards. This is especially true when there is a lot of head motion present, which not only makes it more difficult to correct, but might also lead to measurements which are not sufficiently correctable and have hence to be entirely excluded from further analysis - a situation one would ideally like to prevent. The beneficial effect of any prospective motion correction method will hence be most noticeable in data with substantial head motion. That said, the overall motion of the participants in the current study was rather low, even when the medical tape was not applied. The fact that, despite this, the application of the medical tape led to a significant reduction in head motion, clearly speaks to the efficacy of this motion reduction method, and it is encouraging that this effect was furthermore still traceable in the functional fMRI data. Not only did the application of the medical tape lead to an average increase in tSNR, but it also reduced the amount of motion-related functional connectivity. Furthermore, the amount of this reduction was slightly stronger for connections with larger cortical distances. These observations are in line with previous findings in resting-state data, showing that head motion produces structured noise that causes distance-dependent changes in signal correlations which can bias group results if there are differences in head motion (Power et al., 2012, Power et al., 2014). Eventually, the application of the medical tape also affected task-based fMRI activations in all three cognitive tasks in the tested group of participants, showing significant differences in the estimation of activations in task-relevant brain regions.

Importantly, even though participants in the current study seemed to have been aware of the positive effect the medical tape had on their head motion (as suggested by the questionnaire data), the presence of medical tape did not modulate their behavioural performance in the Stroop task, suggesting that the application of the medical tape did not affect cognitive performance. This is particularly worth noticing since it has been shown that previously suggested motion reduction methods can have an influence on measured variables that are relevant for an empirical study (Yang et al., 2005; Greene et al., 2018).

The current study investigates head motion solely based on estimates that resulted from realignment of the functional MR images. We chose this measure as it is today by far the most common way to quantify MRI head motion. Nevertheless, more sophisticated head position tracking with high speed cameras has recently been made available (Todd et al., 2011; Stucht et al., 2015). Future studies using this technology could provide interesting additional information on the efficacy of the medical tape. In particular, while we show here that the medical tape significantly reduces between-volume head motion, head position tracking with high speed cameras could give insights into whether the medical tape also reduced within-volume motion (a kind of motion that is often ignored in MRI research).

The here tested active head motion reduction method significantly differs from passive methods, such as a plaster cast head holder (Edward et al., 2000) or a bite bar (Bettinardi et al., 1991; Menon et al., 1997). Most notably, the application of the medical tape does *not* aim to fixate participants head and thereby passively prevent them from moving their heads. While the medical tape arguably might put an upper limit on the excess of head motion (as does any form of cushioning, as well as the head-coil and even scanner bore themselves), this upper limit is far away from any head motion that would naturally be expected during an MRI scan session. That is, participants can still visibly move their heads in the range of centimetres under the tape when asked to do so, and any firmer head motion would easily remove the tape entirely (e.g. in case of an emergency). Data from an additional independent participant who was explicitly instructed to move her head confirmed that (a) substantial head motion is still possible when the medical tape is applied and (b) that the head motion observed in the current study was much lower than that. Rather, the medical tape provides tactile feedback by moving the skin on the forehead, making participants aware of their movement and allows them to actively reduce it. Passive fixation furthermore has been reported to be rather unpleasant for the participants (Zaitev et al., 2015). While there is no a priori reason to assume that the application of the medical tape is unpleasant per se, it is nevertheless worth noting that none of the 24 participants in the current study mentioned any discomfort related to the tape, neither during the procedure, nor in the debriefing afterwards. Several participants did, however, positively comment on the usefulness of the feedback information the medical tape provided. The here tested method also significantly differs from another previously presented active head motion reduction method which provides participants with real-time visual head motion information (Yang et al., 2005; Greene et al., 2018). Being a much simpler setup (i.e. no real-time analysis of head motion parameters), implementation is non-technical, quick, cost-efficient and should be applicable in any scanning facility.

Taken together, providing participants with tactile feedback about their head motion by applying medical tape from one side of the MR head-coil, via their forehead, to the other side, is a viable and cost-efficient (both economically and with respect to setup complexity and time investment) method to reduce head motion in MRI in a large variety of scenarios and facilities.

## Acknowledgements

This work was financially supported by Scannexus B.V. as well the European Commission’s Health Cooperation Work Programme of the 7th Framework Programme, under the Grant Agreement n° 602186 (BRAINTRAIN). The authors declare no competing financial interests.

